# Fast Dynamic Whole-Body *In Vivo* Cytometry Using Magnetic Particle Imaging

**DOI:** 10.1101/2025.09.11.675624

**Authors:** Ali Shakeri-Zadeh, Shreyas Kuddannaya, Chengyan Chu, Kritika Sood, Asif Itoo, Cristina Zivko, Vasiliki Mahairaki, Piotr Walczak, J.W.M. Bulte

**Affiliations:** The Russell H. Morgan Department of Radiology and Radiological Science, Division of MR Research, The Johns Hopkins University School of Medicine; Baltimore, MD, USA; Cellular Imaging Section and Vascular Biology Program, Institute for Cell Engineering, The Johns Hopkins University School of Medicine; Baltimore, MD, USA; Center for Advanced Imaging Research, Department of Diagnostic Radiology and Nuclear Medicine, University of Maryland; Baltimore, Maryland, USA; Department of Genetic Medicine, Johns Hopkins University School of Medicine, Baltimore, MD, USA; The Richman Family Precision Medicine Center of Excellence in Alzheimer’s Disease, Johns Hopkins University School of Medicine, Baltimore, MD, USA; Department of Biomedical Engineering, The Johns Hopkins University School of Medicine; Baltimore, MD, USA; Department of Oncology, The Johns Hopkins University School of Medicine; Baltimore, MD, USA; Department of Chemical & Biomolecular Engineering, The Johns Hopkins University Whiting School of Engineering; Baltimore, MD, USA; F.M. Kirby Research Center for Functional Brain Imaging, Kennedy Krieger Institute; Baltimore, MD, USA

## Abstract

Rapid quantification of the immediate organ accumulation of injected stem cells remains a major challenge. We performed *in vivo* cytometry (non-invasive cell counting) with magnetic particle imaging (MPI) to track magnetically labeled cells in real-time on a time scale of minutes with high sensitivity, zero background signal, and simple linear quantification. Human mesenchymal stem cells (hMSCs, ∼25 µm in diameter) and human neural precursor cells (hNPCs, ∼10 µm in diameter) were labeled with ferucarbotran or Synomag®-D70 as superparamagnetic iron oxide (SPIOs), and tracked with MPI in mice to map their whole-body cell biodistribution after intravenous (IV) or intra-arterial (IA) injection. The organ site of cell accumulation and retention were dependent on cell type, injection route, and frequency of administration, with the lung and liver acting as the major entrapment organs. *In vivo* MPI enabled quantitative tracking of the dynamic clearance and redistribution of labeled cells, showing major differences between larger hMSCs and smaller hNPCs. Co-registered MRI/CT and histological validation confirmed the anatomical localization of SPIO-labeled cells including the brain following IA injection. Integrating MPI cytometry with preclinical and translational studies may aid in further optimization of the route, dose, and frequency of stem cell administration.

**One Sentence Summary:** Fast whole-body *in vivo* cytometry using MPI is able to dynamically track and quantify therapeutic stem cell accumulation.

## INTRODUCTION

Developing methods to track and quantify therapeutic stem cells and immune cells *in vivo* remains an active area of research to successfully translate cell therapies into clinical practice (*1*). It has been shown that cell injections can miss their target in up to 50% of patients even when injected under ultrasound imaging-guided injections (*2*), emphasizing the need for more effective methods to monitor and improve cell delivery (*3*). Traditional imaging modalities for *in vivo* cell tracking often fall short in providing the necessary sensitivity, specificity and quantification required for precise whole-body cell tracking. ^1^H magnetic resonance imaging (MRI) has limited specificity with poor quantification to detect and quantify magnetically labeled cells (*4, 5*). Nuclear imaging, either positron emission tomography (PET) or single photon emission computed tomography (SPECT), is a quantitative method and offers high sensitivity for cell tracking but involves ionizing radiation and lacks spatial resolution (*6*). Optical imaging suffers from limited tissue penetration and quantification difficulties (*7*). Magnetic particle imaging (MPI) has emerged as a promising modality for *in vivo* cell tracking (*8-11*) with a reported *in vivo* detection sensitivity as low as 250 labeled cells (*12*). MPI directly detects superparamagnetic iron oxide (SPIO) nanoparticles with high sensitivity and specificity (*13, 14*). Importantly, MPI provides a linear relationship between signal intensity and SPIO concentration, enabling accurate quantification of SPIO-labeled cells and tracers (*8, 10, 15*). ^19^F MRI has been explored as an alternative quantitative cell tracking approach for *in vivo* cytometry (*16*), but it suffers from lower sensitivity and temporal resolution compared to MPI (*17*). The main limitations of MPI include its relatively low spatial resolution and its lack of anatomical context, necessitating its combination with imaging modalities such as MRI or computed tomography (CT) for accurate anatomical localization.

Recent progress in the development of human-scale MPI scanners demonstrates that the technology is scalable and positioned for clinical translation (*18-21*).The use of nonionizing magnetic fields in MPI (*22*), combined with the biocompatibility and minimal functional impact of the magnetic labeling of cells (*23*), supports the future use of clinical MPI for real-time cell tracking.

Here, we introduce the concept of *in vivo* MPI cytometry, an imaging approach that enables dynamic whole-body tracking and quantification of SPIO-labeled therapeutic cells in living subjects on a time scale of minutes. By implementing fiducial-based calibration standards and optimized MPI/MRI/CT imaging protocols, this method allows accurate cell quantification across multiple organs over time, offering a series of comprehensive snapshots of cell biodistribution. I*n vivo* MPI cytometry was performed on two distinct stem cell types, i.e., human mesenchymal stem cells (hMSCs), which are relatively large cells (∼25 µm in diameter) used before for MPI cell tracking studies (*8, 24*), and human neural precursor cells (hNPCs), representing smaller cells (∼10 µm in diameter) used with MRI cell tracking (*25*). Our results reveal distinct biodistribution profiles for the two cell types, highlighting the suitability of MPI cytometry for quantitative monitoring of cell distribution *in vivo*. Although previous studies have used MPI to track cells *in vivo*, cells were detected at a single implantation site or at a single end time point.

## RESULTS

### Whole-body quantitative *in vivo* MPI cell tracking

hMSCs were efficiently labeled with ferucarbotran (**Fig. 1A**). *In vivo* MPI/CT imaging revealed distinctly different biodistribution profiles for IV compared to IA injection (**Fig. 1B,C**). At 30 min post-IV injection cells accumulated within the lungs, with negligible signal in liver and brain, indicating pulmonary entrapment. In contrast, IA injection revealed cell localization in the brain, lungs, and liver. After 1 day, IV-injected cells completely redistributed to the liver, while IA-injected cells persisted in both brain and liver without lung signal (**Fig. 1B**). Using a calibration curve for the same batch of labeled hMSCs that were injected, the number of cells were at 30 min post-IA injection were calculated to be 99,937±10,392 for the lung/liver and 27,500±2,240 cells for the brain. These total numbers summing up to approximately 125,000 cells closely equals the amount of 1.2×10^5^ injected cells, translating to an iron mass of 57 pg Fe per cell as estimated from signal intensity and known Fe content determined using a Ferrozine assay.

**Figure 1.**
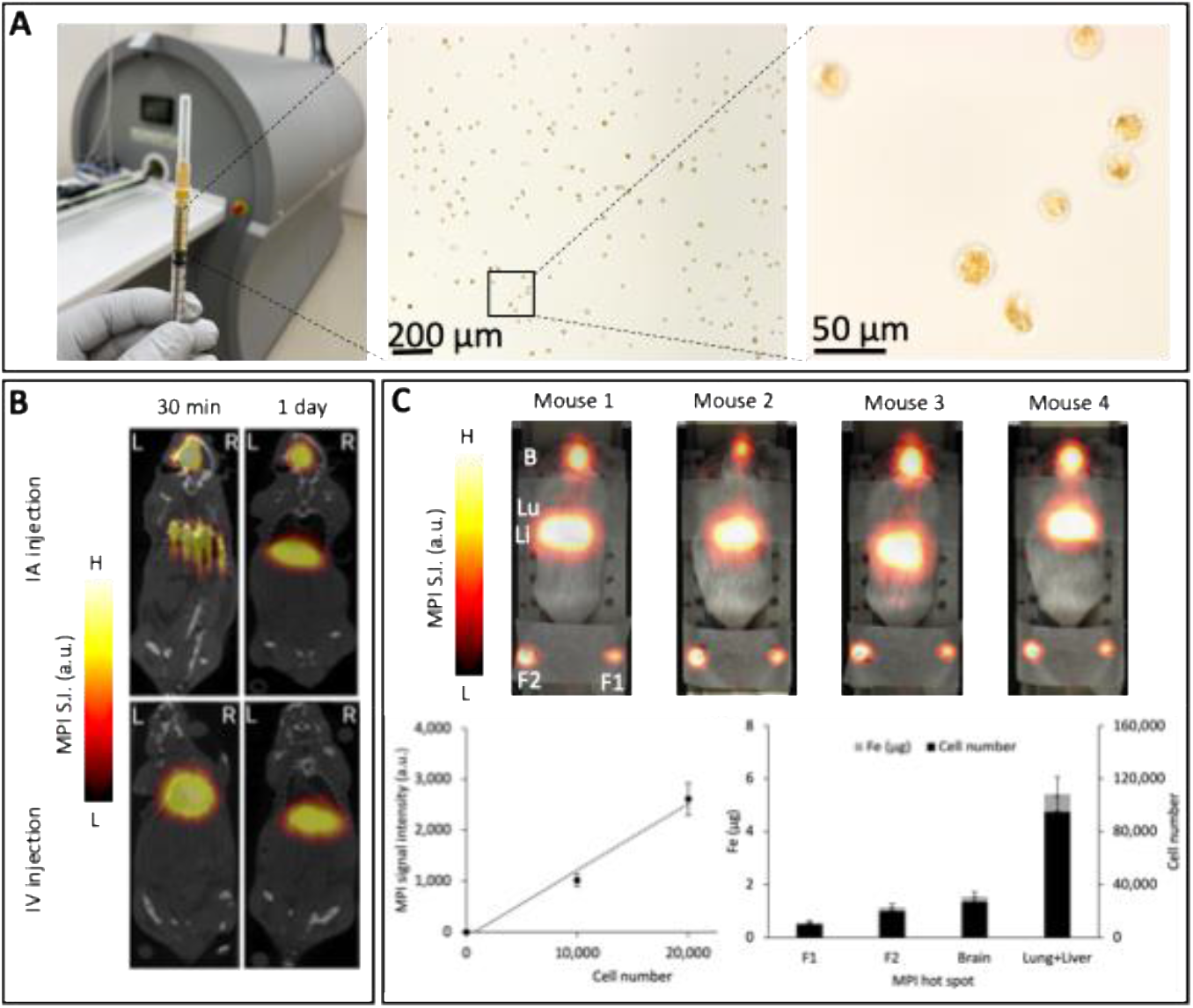
Whole-body *in vivo* tracking and quantification of hMSCs following IV or IA injection. (**A**) Macro- and microscopic visualization of ferucarbotran-labeled hMSCs before injection. (**B**) *In vivo* MPI/CT images of labeled hMSCs at 30 min and 1 day after IV or IA injection. (**C**) Quantitative analysis of cell accumulation 30 min post-IA injection. Two fiducials (F1=1×10^4^ and F2=2×10^4^ labeled cells) were used for calibration (y = 0.13x - 94.92; R^2^ = 0.98). Data in each figure panel represent three independent experiments. B, Lu, Li, and F stand for brain, lungs, liver and fiducials, respectively.

### Bimodal MPI/MRI cell tracking

To validate the cerebral delivery of ferucarbotran-labeled hMSCs following IA injection, we performed *ex vivo* magnetic imaging (**Fig. 2A**) and histological analysis (**Fig. 2B**). Coronal MRI scans showed unilateral hypointense regions across multiple brain slices, consistent with labeled cell accumulation. The signal voids were present from midbrain to forebrain, indicating a global cell distribution in the entire ipsilateral hemisphere. MPI scans of the same brains revealed hot spots that overlapped with the MRI hypointensities. Histological Prussian blue staining further confirmed the presence of SPIO-labeled hMSCs. Anti-human nuclear antigen (HuNa) staining revealed that the iron-loaded cells contained human nuclei, confirming they were indeed the hMSCs that were injected, corroborating the MRI/MPI findings.

**Figure 2.**
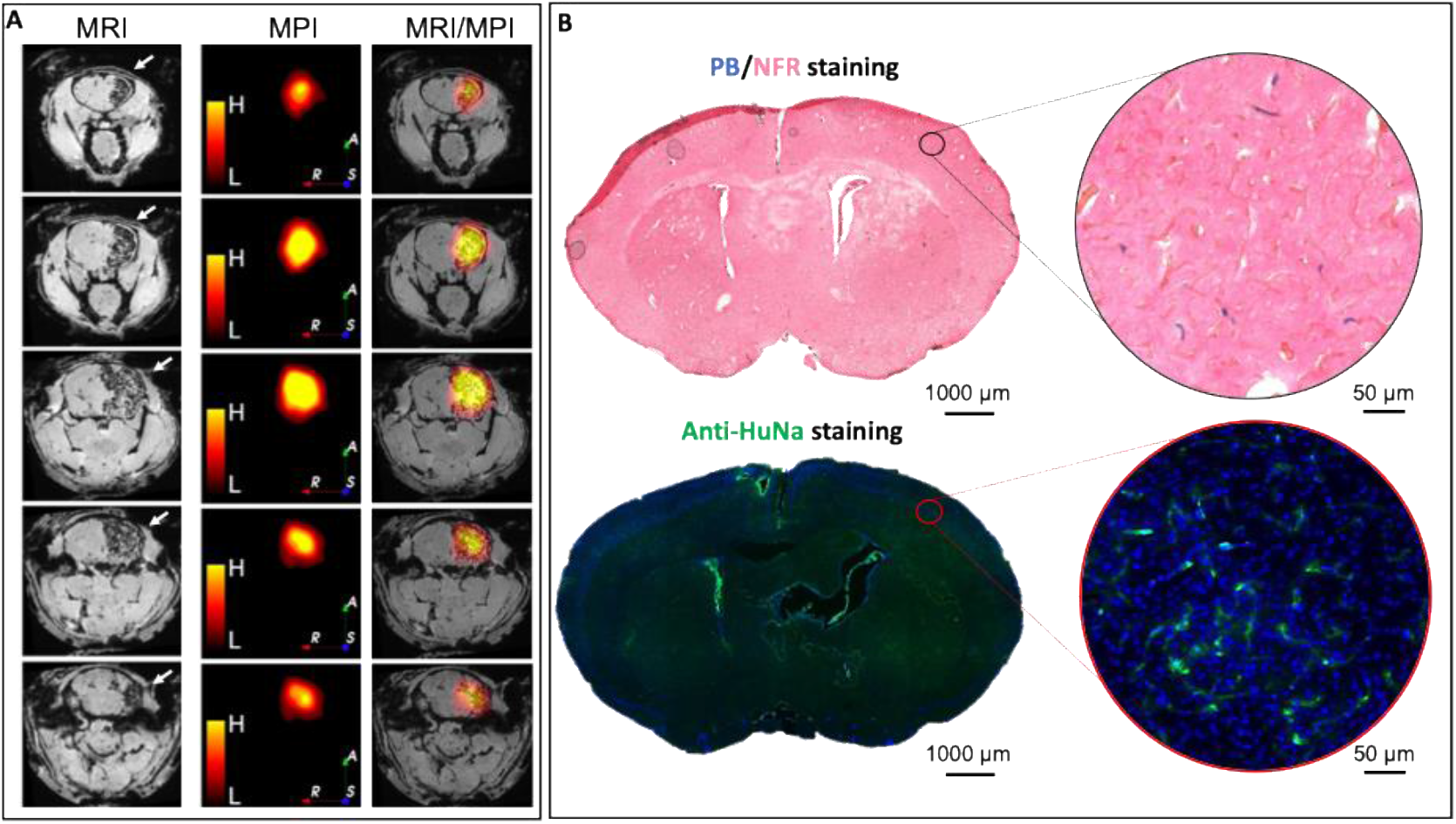
Bimodal MPI/MRI of hMSCs at 6 hours after IA injection. (**A**) Consecutive coronal *ex vivo* MRI, MPI, and MRI/MPI overlay images showing MRI hypointensities (white arrows) and MPI hot spots from midbrain to forebrain, corresponding to labeled hMSCs. (**B**) Corresponding coronal brain sections stained for iron (top row) and anti-HuNa (bottom row), demonstrating that the remains localized in hMSCs.

### Longitudinal MPI cell tracking

To investigate the long-term biodistribution and persistence of IA-injected cells, we performed serial *in vivo* and *ex vivo* MPI scans at 6 h (day 0), 2 days, 7 days, and 30 days post-injection of 120,000 SPIO-labeled hMSCs (**Fig. 3A**). Using two fiducials containing labeled hMSCs from the same batch that was injected, MPI signal quantification revealed 22,294±1,920 cells in the brain and 114,940±13,125 cells in the lungs/liver on day 0 (**Fig. 3B**). By day 2, the brain signal became undetectable *in vivo*, indicating that most cells redistributed from the brain to other organs. The lungs and liver retained 117,385±14,450 cells at Day 2, reflecting the early loss of brain-localized cells and sustained hepatic/pulmonary retention. The cell number in liver declined to 58,835±5,210 at day 7 and 24,280±2,225 at day 30. *Ex vivo* MPI confirmed these observations with no detectable cells in the heart. At Day 0, 8,200±970 cells were detected in the brain, 5,430±436 in the lungs, and 102,500±11,240 in the liver, providing a (lung+liver)/brain ratio of ∼13. By day 2, only 2,330±190 cells remained in the brain and no cells were detected in the lungs, while the liver signal increased to 108,800±8,425 cells, yielding a (lung+liver)/brain ratio of ∼47. At days 7 and, the liver retained 67,300±7,360 and 22,240±2,015 cells, respectively, with no detectable cells in the lungs or brain. These findings suggest a possible redistribution of SPIO-labeled cells from the lungs and brain into the liver between days 0 and 2, followed by gradual clearance.

**Figure 3.**
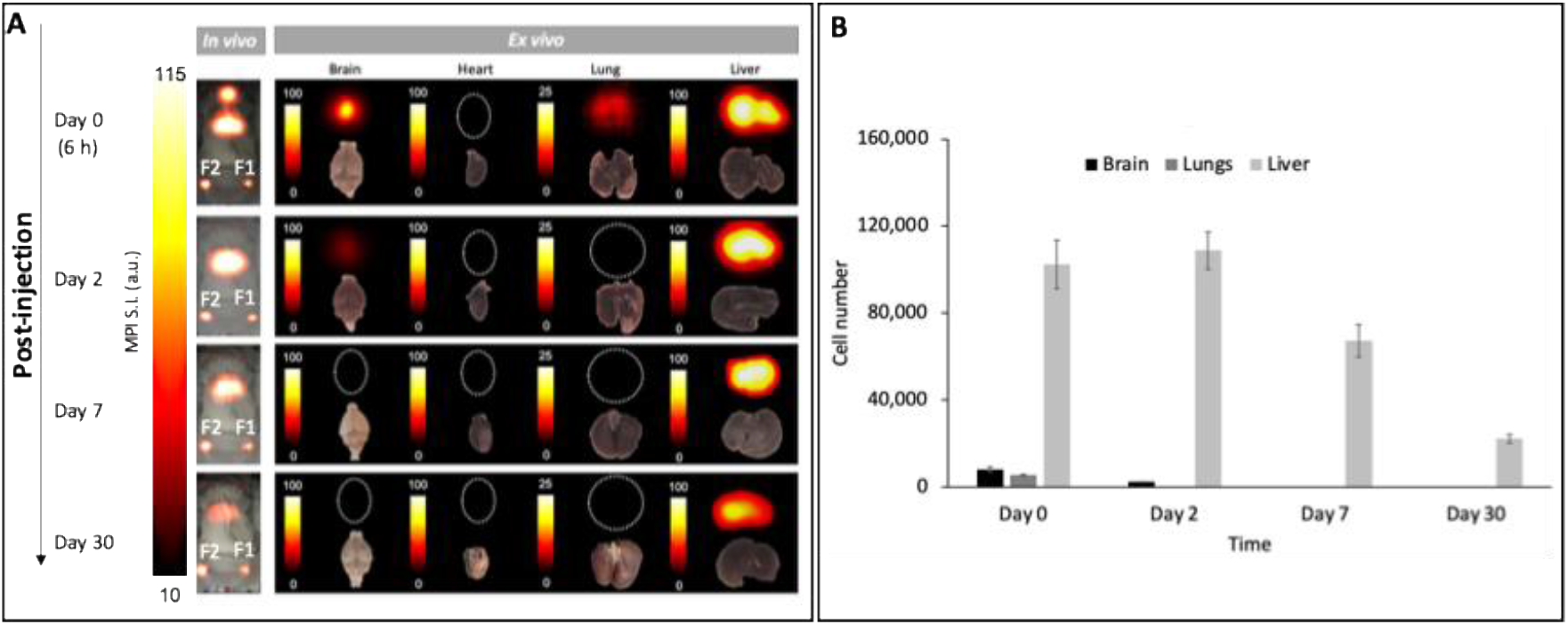
Longitudinal MPI of labeled hMSCs after IA injection. (**A**) In vivo and ex vivo MPI at different time points post-injection. Color-coded heat maps indicate signal intensity. Dotted circles highlight organs with no detectable signal. Shown are representative data from three independent experiments. (**B**) *Ex vivo* MPI-based quantification of labeled hMSCs. Similar to the *in vivo* studies, F1=1×10^4^ and F2=2×10^4^ labeled cells were used as fiducials for *ex vivo* quantification.

### Fast dynamic whole-body *in vivo* MPI cytometry: effect of cell size

To compare the *in vivo* biodistribution of two stem cell types having different sizes, we performed dynamic MPI tracking of 120,000 IA-injected ferucarbotran-labeled hMSCs and 960,000 Synomag®-D70-labeled human neural precursor cells (NPCs), have spherical diameters of ∼25 and 10 µm following trypsinization, respectively (**Fig. 4**). Each total cell dose was injected using four incremental injections, with equal cell numbers delivered in each injection. Both cell types were effectively labeled as confirmed with PB staining. For hMSCs serial *in vivo* MPI performed over 12 minutes post-injection revealed a prominent signal in both the brain and the liver/lung regions, with the signal increasing over time from 0 to 12 min. Using a calibration curve obtained from fiducials containing known numbers of labeled cells from the same batch as was injected, it could be calculated that 22,569±1,805 hMSCs localized in the brain, with 112,808±10,600 cells in the combined liver and lung regions, yielding a (lung+liver)/brain ratio of ∼5. The labeled hMSCs were also quantified using *ex vivo* MPI (**Fig. 5A**), which detected 10,614±1,004 cells in the brain, 7,212±820 in the lungs, and 92,140±10,437 in the liver. The total calculated distribution numbers of 135,000 (*in vivo* MPI) and 110,000 (*ex vivo* MPI) are in close agreement with the total amount of injected cells, being 120,000 (**Table 1**).

**Table 1.** *In vivo* and *ex vivo* MPI cytometry of SPIO-labeled hMSCs and hNPCs.

| Parameter | hMSCs | hNPCs |
| --- | --- | --- |
| Injected cell dose | 120,000 | 960,000 |
| Cell size (μm) | ~25 | ~10 |
| **In vivo MPI (12 min post injection)** |  |  |
| Brain (mean ± SD) | 22,569 ± 1,805 | 74,240 ± 6,854 |
| Lung + Liver (mean ± SD) | 112,808 ± 10,600 | 474,580 ± 39,094 |
| Total detected cells | 135,377 | 548,820 |
| % of injected cells detected | 113% | 57.2% |
| **Ex vivo MPI (6 hours post injection)** |  |  |
| Brain (mean ± SD) | 10,614 ± 1,004 | 3,420 ± 280 |
| Lungs (mean ± SD) | 7,212 ± 820 | Not detected |
| Liver (mean ± SD) | 92,140 ± 10,437 | 143,540 ± 12,430 |
| Lung + Liver (mean ± SD) | 99,352 ± 10,719 | 143,540 ± 12,430 |
| Total detected cells | 109,966 | 146,960 |
| % of injected cells detected | 91.6% | 15.3% |

**Figure 4.**
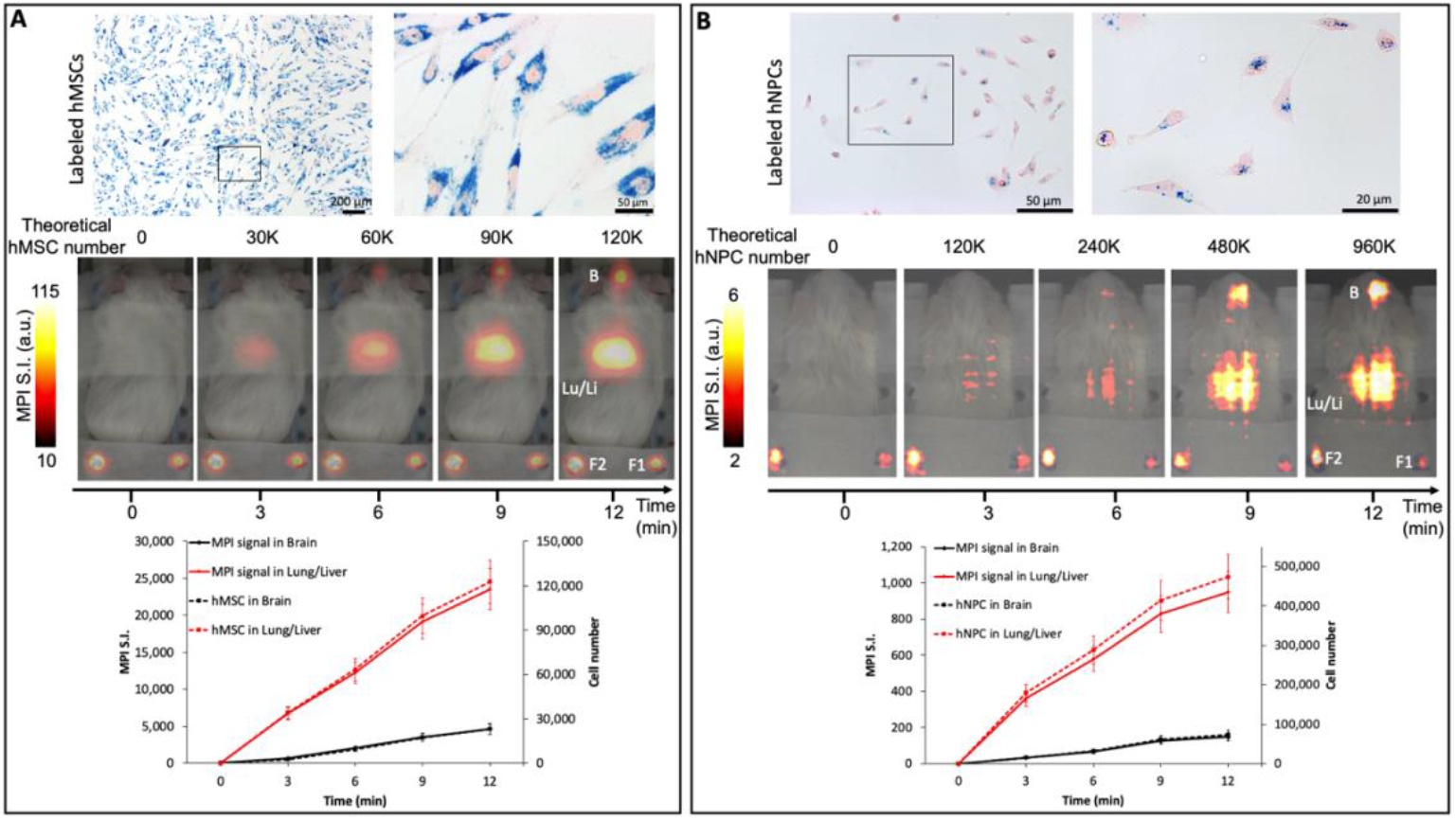
Dynamic *in vivo* MPI cytometry comparison of labeled hMSCs (A) and hNPCs (B) after IA injection. **Top panels** show PB staining to confirm intracellular iron accumulation in each cell type. **Middle panels** display time-lapse MPI scans at multiple time points post-injection. B = brain, Lu/Li = lung/liver. Fiducials F1 and F2 contain 10,000 and 20,000 labeled hMSCs in panel (A), and 20,000 and 40,000 labeled hNPCs in panel (B), respectively, from the same batch of injected cells. The calibration equations used were: *y* = 0.1868*x* + 220 (R^2^ = 0.99) for hMSCs and *y* = 0.0021*x* – 2 (R^2^ = 0.99) for hNPCs. **Bottom panels** present quantitative plots of MPI signal intensity (arbitrary units) and calculated cell numbers over time in each region, demonstrating the ability of MPI for both fast localization and dynamic quantification of labeled cells. Data represent three independent experiments.

**Figure 5.**
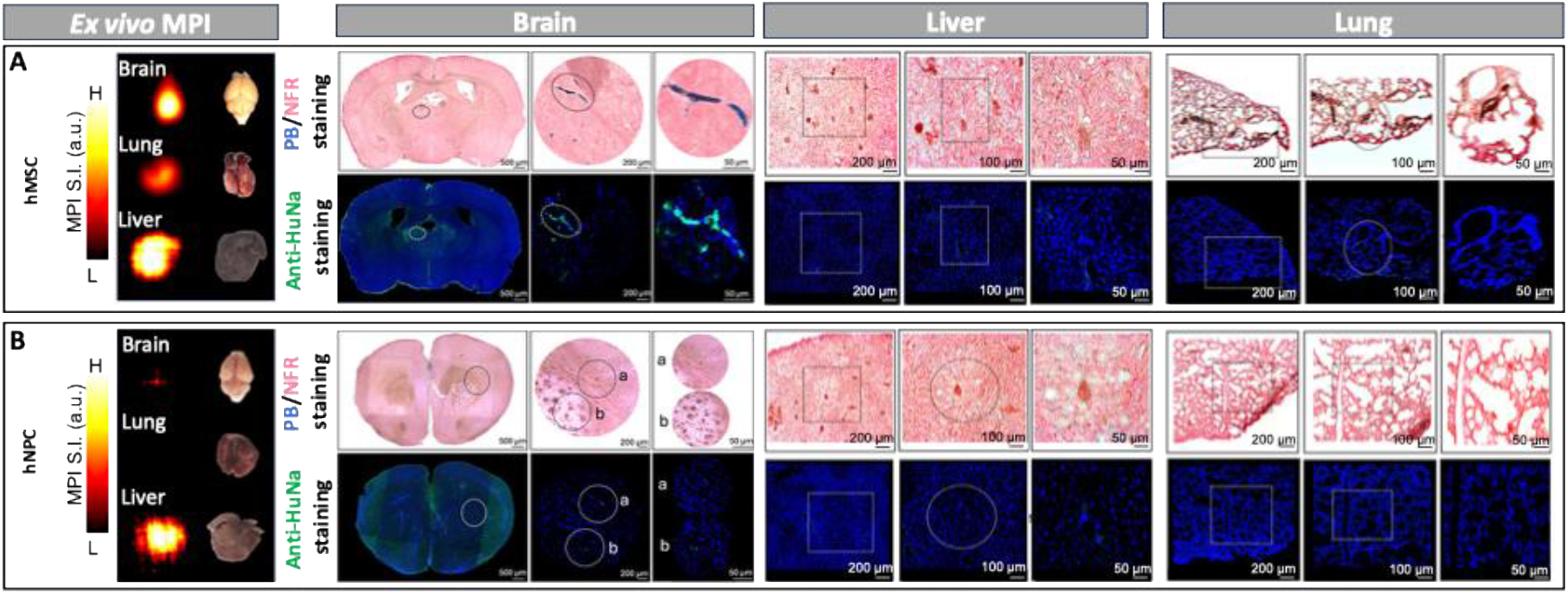
*Ex vivo* MPI and histology validate cell localization in major organs. Representative *ex vivo* MPI and histological analysis of ferucarbotran-labeled hMSCs (**A**) and SynomagD-70-labeled hNPCs (**B**) 6 hours after IA-injection. Co-localization of positive PB and anti-HuNA staining confirms the presence of SPIO-labeled hMSCs in the brain.

For the smaller hNPCs (**Fig. 4B**), the *in vivo* MPI demonstrated a dynamic signal evolvement in the same anatomical regions. At 12 min post injection, ∼57% of injected hNPCs were detected *in vivo*, including 74,240±6,854 hNPCs in the brain and 474,580±39,094 cells in the lung and liver, corresponding to a (lung+liver)/brain ratio of approximately ∼6.5, i.e. ∼30% higher than the corresponding ratio observed for hMSCs. The ∼43% discrepancy between the injected and detected hNPCs may be attributed to several factors, including diffuse biodistribution below the MPI detection threshold and the inherently lower iron loading of hNPCs relative to hMSCs. Additionally, limited spatial resolution and spillover effect in MPI, particularly from high-signal regions such as the liver, may obscure weak or widely dispersed signals, further contributing to underestimation. *Ex vivo* MPI (**Fig. 5B**) detected only 3,420 ± 280 cells in the brain, no cells in the lungs, and 143,540 ± 12,430 cells in the liver (**Table 1**). Histological analysis with PB staining and anti-HuNa immunolabeling only confirmed the presence of SPIO-labeled human cells in the hMSC-injected brains. As hNPCs are much smaller than hMSCs this could have resulted from a larger wash-out during post-mortem processing compared to hMSCs. Although MPI revealed substantial accumulation of labeled hMSCs in the liver and lungs as well as accumulation of labeled hNPCs in the liver, histological analyses failed to corroborate the corresponding MPI signals. This apparent discrepancy may be attributed to multiple technical and biological factors. First, MPI offers high sensitivity and full-organ detection capability, while histology is inherently limited to thin tissue sections and may miss sparsely distributed cell populations. Second, cells trapped in the microvasculature, especially within the brain, may be lost during organ removal and perfusion, leading to false negatives in post-mortem histological analysis. Similarly, in the liver and lungs, cells may localize transiently within sinusoidal spaces or vascular compartments and be washed out during fixation and sectioning.

## DISCUSSION

### Route of delivery: IA vs. IV cell injection

The findings in this study show how the route of cell delivery largely influences the initial biodistribution and subsequent homing. IA injection of SPIO-labeled cells led to a considerably different organ distribution compared to IV injection. Consistent with numerous prior imaging studies, IV-injected hMSCs underwent a first-pass sequestration in the lung vasculature (*24, 26*). This well-known “pulmonary trap” is attributed to the large size (18–30 μm diameter) and adhesive properties of hMSCs, which cause 50–80% of IV injected cells to lodge in the pulmonary capillaries (*26*). Our MPI data are in alignment as we observed a strong MPI signal concentrated in the lungs 30 min after IV injection (**Fig. 1B**). This mirrors previous reports using both optical and nuclear imaging. Bioluminescence imaging of luciferase-expressing hMSCs shows a bright lung signal within minutes of IV injection, which diminishes over 1–2 days as cells clear or die (*26*). Similarly, SPECT and PET tracking of radiolabeled hMSCs have documented that IV injected cells accumulate first in lungs (*26, 27*), with a subsequent redistribution in the liver (*28*). Consequently, only a small fraction of hMSCs can reach certain targeted tissues after IV injection. Delivering cells via a targeted arterial route allows a greater proportion of injected cells to reach the intended organ. In our study, IA injection greatly enhanced cell delivery to the brain, whereas an IV injection with an ∼8-fold higher dose yielded no detectable brain MPI signal. This observation aligns with prior reports using other modalities. T2-weighted MRI demonstrated that IA transplantation of hMSCs led to markedly enhanced cell homing in the brain of stroked rats, while IV injection of the same cells produced negligible brain localization (*29*). However, IA injection of hMSCs may carry a risk of microvascular blockage (*29*) and hence its safety must be carefully considered in terms of cell size and cell dose (*30*). Nevertheless, the ability of IA administration to bypass the lung “barrier” and achieve near whole-body distribution is a clear advantage for targeting organs beyond the pulmonary circuit, as demonstrated for targeted renal delivery of hMSCs to the kidney using the renal artery (*31*), or bone marrow stem cells to the pancreas and spleen using the superior pancreaticoduodenal artery or splenic artery, respectively (*32*).

### Cell size differences: hMSCs vs. hNPCs

Beyond the delivery route, the intrinsic properties of the cells led to marked differences in biodistribution and organ retainment (**Fig. 4**). A robust brain signal could only be observed after injection of 480,000 hNPCs. One key factor which plays a major role in the kinetics of cell type– dependent biodistribution is cell size. The hMSCs used here average ∼25 μm in diameter, with reported ranges up to 31 μm (*33*), roughly double the size of hNPCs, whose cell bodies are on the order of 9–12 μm. This size difference underlies the reduced cerebral trapping we observed with hNPCs compared to hMSCs (**Figs. 4** and **5**). Additionally, differences in cellular surface markers and adhesion molecules between hMSCs and hNPCs could play a role. MSCs express high levels of integrins and other receptors that facilitate adhesion to endothelial cells (*34*). NPCs, being neural-lineage cells, may have a different adhesion molecule profile and may exhibit less “stickiness” for blood vessels.

### MPI vs. other conventional tracking modalities

MPI enables a level of cell tracking sensitivity and quantification that addresses many limitations of established imaging modalities (MRI, PET/SPECT, and optical imaging). A major advantage of MPI is the ability to quantitatively track small cell populations in deep tissue with high specificity (**Fig. 2**), as there is no background signal. Hence, there is no ambiguity in identifying SPIO-labeled cells (*14, 24*).This specificity contrasts with conventional MRI of SPIO-labeled cells, which relies on negative contrast that can be confounded by endogenous sources (e.g. air/tissue interfaces, hemorrhage, or calcifications). Hence, linear quantitation of iron using MPI enabled us to convert image signal directly into cell number estimates, which is not feasible with MRI. As a result, we could “non-invasively” count the number of cells, i.e., perform “in vivo cytometry” in each organ over time. We are able to detect and quantify cell numbers in the order of thousands with MPI *in vivo*, which is in line with published sensitivity benchmarks. A recent direct comparison showed that MPI can detect as few as 4,000 hMSCs (approximately 75 ng of iron) in a mouse, whereas ^19^F MRI may be required on the order of 250,000 cells for detection (*17*).

Another remarkable advantage of MPI is its capacity for its dynamic imaging. We leveraged fast scanning and the lack of tissue attenuation in MPI to monitor cell biodistribution immediately after injection and then repeatedly over days to weeks. Such data are difficult to obtain with other modalities. Optical imaging cannot be used in deep organs with tissue attenuation, while ^19^F MRI requires lengthy scans. Acquiring just eight contiguous 5.0 × 2.8 cm (2-mm thick) slices with ^19^F MRI takes approximately 2 hours, without whole-body coverage (*16*). MPI enables rapid whole-body imaging of mice, requiring only 1–2 min for 2D scans, depending on imaging parameters such as scan mode. Recent work has demonstrated real-time “fluoroscopy” with MPI, tracking labeled cells moving through a flow system at sub-second time frame rates (*35*). In our studies we did not require sub-second time frames, but we did acquire serial whole-body snap shots immediately post-injection to observe the first-pass distribution. The ability to see cells in near real-time opens the door to optimizing injection techniques (e.g., slow vs. bolus injection, use of magnetic targeting, etc.) in ways that static endpoint measurements cannot.

Although MPI currently achieves a practical spatial resolution of approximately 1 mm (*36*), which can be enhanced to ≤300 μm with optimized SPIO tracers and pulse sequences (*37*), its current resolution remains comparable to that of preclinical PET (∼1–2 mm) (*38*) and ^19^F MRI (∼0.8 mm) (*16*). However, it still lags behind the sub-100 µm resolution achievable with high-field conventional MRI. Limited spatial resolution of MPI can be effectively mitigated by co-registering MPI with high-resolution MRI (**Fig. 2A**), allowing the superimposition of quantitative MPI-based cell tracking onto detailed anatomical maps for accurate and spatially resolved cytometry.

### Clinical implications and translational outlook

Our study demonstrates a tracking platform with clear translational potential for integration with (stem) cell therapies. First, the tracer we employed (SPIO) is intrinsically suitable for human use. SPIO agents such as ferucarbotran, formerly used as the MRI contrast agent Resovist®, and now re-introduced as Resotran® (*39*), have well-documented safety profiles in patients, and they can be used to label therapeutic cells without impairing their function (*23*). The concentration of iron per cell achieved here (tens of pg Fe/cell) is on par with that used in prior human MRI cell-tracking trials indicating that our labeling approach is scalable to clinical cell doses. Moreover, MPI itself is advancing toward clinical utilization. MPI uses low-frequency magnetic fields that are far below FDA limits and pose no known risk to tissues (*40*). Recent engineering breakthroughs have produced human-scale MPI scanners (*18-21*), and first-in-man studies are now being conducted. As a result, the combination of a non-toxic clinically approved tracer and a safe imaging modality positions MPI-based cell tracking well for rapid clinical translation. MPI *in vivo* cytometry may address important questions such as if the cells reach the target organ, when, how many, and do they persist or are rapidly cleared.

## MATERIALS AND METHODS

### Study design

This study was designed to evaluate the feasibility, reproducibility and quantitative capability of *in vivo* MPI cytometry for whole-body tracking of two SPIO-labeled stem cell types (hMSCs and hNPCs) having different sizes. Cells were injected into immunodeficient Rag2 mice using either IA or IV injection. Dynamic and longitudinal MPI scans were performed to assess whole-body distribution and organ-level cell quantification, using fiducial-based calibration for standardization. MRI or CT was used for anatomical referencing, and the imaging data were verified using histological assessment.

### hMSC labeling

P2 human bone marrow-derived MSCs were obtained from RoosterBio™ and expanded in culture up to P5 using MSC basal medium (MSCBM, Lonza) supplemented with 10% MSC growth supplement, 2% l-glutamine, 0.1% gentamicin, and 0.1% amphotericin. hMSCs were incubated for 24 h with a pre-prepared labeling medium consisting of MSCBM medium supplemented with ferucarbotran (25 µg Fe/ml, Meito Sangyo Co.) and poly-L-lysine (Sigma P-1524, 3 µg/ml). Following incubation, the labeling medium was removed, and the labeled hMSCs were washed three times with PBS to remove unbound particles.

### hNPC labeling

Induced pluripotent stem cells (iPSCs) derived from a patient with amyotrophic lateral sclerosis and genetically corrected for the SOD1 mutation (39B2.5 line) were obtained from Dr. Kevin Eggan (*41*). iPSCs were differentiated into NPCs using a defined neural induction medium supplemented with dual-SMAD inhibitors (*42*). Briefly, iPSCs were plated on matrigel-coated plates and cultured to confluence in Essential 8 medium (Gibco). Neural induction was initiated using WiCell neural induction medium supplemented with dual-SMAD pathway inhibitors including LDN193189 (Stemgent) and SB431542 (Sigma). Starting on day 3, retinoic acid (Sigma) was added to promote caudalization of the neuroectoderm. On day 5, the medium was replaced with neural induction medium containing retinoic acid and continued until day 12. From day 12 to day 25, cells were maintained in neural induction medium supplemented with 1 μM purmorphamine (Sigma), 10 ng/ml BDNF (Invitrogen), 10 ng/ml GDNF (R&D Systems), 10 ng/ml CNTF (R&D Systems), 200 µM ascorbic acid (Sigma) and 10 ng/ml IGF-1 (R&D Systems) to support NPC maturation. Cells were harvested on day 25 for subsequent labeling with Synomag®-D70.

For hNPCs, the labeling procedure was performed in a standard 24-well tissue culture plate. Each well contained 100,000 hNPCs suspended in 2 ml of complete growth medium. Cells were labeled with Synomag®-D70 (40 µg Fe/ml), and poly-L-lysine (390 ng/ml). Cells were incubated for 4 h at 37°C. After incubation, cells were washed three times with 10 mM phosphate buffered saline (PBS, pH =7.2-7.4).

### Prussian blue staining and Ferrozine iron assay

PB staining was performed to detect the presence of iron within labeled cells. Cells were fixed with 4% glutaraldehyde for 20 mins, washed with PBS, and incubated with Perl’s reagent (potassium ferrocyanide in HCl) for 30 min at room temperature. To enhance cellular visualization, cells were counterstained with nuclear fast red (NFR) for 20 mins. Stained cells were washed with deionized water to remove excess reagent and then imaged using a Zeiss Apotome 2 microscope. The Fe content in labeled cells was determined using a Ferrozine assay as described elsewhere (*43*). A standard curve prepared with known Fe concentrations was used to calculate the Fe content in the sample.

### *In vivo* injection procedures

All animal procedures were approved by our Institutional Animal Care and Use Committee. Male immunodeficient Rag2^−/−^ mice (6–8 weeks old, Jackson Laboratory) were housed under a 12-hour light/dark cycle with ad libitum access to food and water. All injection, surgery and *in vivo* imaging procedures were performed under anesthesia using 5% induction and 1.5% maintenance isoflurane. Before injection, all labeled cell suspensions were assessed for iron content and labeling efficiency. For IA injection, the ICA was surgically catheterized as previously described (*30, 44*) and cells were delivered using a Harvard Apparatus infusion pump. IV injections were performed via a tail vein catheter and a 27G needle using the same pump system. Cells were suspended in 200 μl of sterile PBS and delivered either as a single injection or in four sequential injections at 3-minute intervals for dynamic imaging studies. The infusion rate was maintained at 0.15 mL/min. Following IA injection, animals received a single subcutaneous dose of buprenorphine ER (1 mg/kg) for postoperative analgesia.

### MPI

MPI was performed using a Momentum field-free line scanner (Magnetic Insight Inc.). A custom mouse bed with two lateral racks, each featuring holes to hold fiducials, was designed in FreeCAD (v0.21.0) and fabricated using a 3D printer (Ultimaker 2 Extended+). For *in vivo* imaging, a 2D whole-body MPI scan was first acquired in standard mode, followed by a 3D scan (also in standard mode) using 21 projections. The standard mode was selected for its rapid acquisition speed, making it well suited for dynamic *in vivo* imaging (*45*). Prior to use, the imaging bed was scanned to confirm the absence of magnetic contaminants. Two fiducials containing different numbers of SPIO-labeled cells (from the same batch as those injected) suspended in 100 µL PBS were positioned within the field of view (FOV) for both MPI signal quantification and image co-registration. Regions of interest (ROIs) were manually delineated on “sharpened” MPI images using ImageJ. Windowing parameters (brightness, contrast, minimum, and maximum) were consistently adjusted across datasets acquired under the same conditions to minimize blooming artifacts, particularly in areas with high signal intensity (e.g., liver). MPI signal intensity for each hot spot was calculated by multiplying the mean signal intensity by the ROI area.

### Micro-CT

Using the same *in vivo* MPI bed equipped with a custom 3D-printed adaptor for co-registration, mice were maintained under anesthesia and promptly transferred to an adjacent IVIS Spectrum/CT system (Caliper Sciences) to acquire anatomical micro-CT images. Micro-CT scans were performed with a tube voltage of 50 kVp, a current of 200 µA, and an exposure time of 300 ms. Images were acquired at a voxel resolution of 100 µm, and 3D reconstructions were generated using 3D Slicer software.

### MRI

High-resolution MRI was performed *ex vivo* only. To enable co-registration of MPI and MRI data for the mouse head, a custom MPI/MRI cradle was designed using FreeCAD (v0.21.0) and fabricated with a 3D printer (Ultimaker 2 Extended+). The cradle included two alignment disks, each with a hole to hold fiducial tubes for spatial referencing. Following MPI acquisition of the excised mouse head mounted in a 3D-printed cradle, high-resolution *ex vivo* MRI was performed using a Bruker BioSpec 17.6T vertical bore scanner equipped with a Bruker BioSpin 25-mm proton transmit/receive volume coil and ParaVision 5.1 software. A 3D FLASH gradient-echo sequence was used with the following parameters: echo time=2.6 ms, repetition time=8.4 ms, flip angle=5°, FOV=20×30×18 mm^3^, and matrix size=200×300×180, resulting in an isotropic spatial resolution of 100 µm. To improve the signal-to-noise ratio, 16 signal averages were acquired.

### MPI/MRI and MPI/CT data co-registration

Co-registration was performed using the Volume, Look-Up Table, Transform, and Rendering tools in 3D Slicer. Two fiducial tubes containing labeled cells were included in the imaging setup and used as reference points for accurate alignment of MPI and MRI/CT datasets.

### Histopathology

Following *in vivo* imaging, mice were sacrificed and transcardially perfused first with 1x PBS, followed by 4% paraformaldehyde before the brain, lungs, heart and liver were collected for *ex vivo* imaging and post-mortem analysis. Tissues were immersed in 4% PFA at 4 °C for 24 h, and then transferred to 30% sucrose for another 72 h. After embedding in optimal cutting temperature compound, the organs were sequentially cut into 20 μm sections using a cryostat (Thermo Fisher Scientific). For PB staining, tissues were fixed with 4% glutaraldehyde for 20 min and stained as described above. Anti-HuNa staining was performed to visualize hMSCs or hNPCs. Tissue sections were first blocked with 5% BSA and 0.1% Triton X-100 in Tris-buffered saline (TBS, pH=7.0) for 2 h, followed by incubation with mouse anti-HuNa (1:250, NBP2-34342, Novus-Bio). Goat anti-mouse Alexa-fluor 488 (1:500, A-11001, Invitrogen) was used as secondary antibody, prepared in 3% BSA in TBS. Sections were incubated for 2 h at room temperature and then triple washed in 1×TBS to remove unbound antibody in each step. Sections were cover-slipped with mounting medium containing DAPI. Fluorescence microscopy was performed using a Zeiss Axiovert 200 M inverted epifluorescence microscope.

### Statistical analysis

Linear regression analysis was performed to plot the total MPI signal (a.u.) for each fiducial against Fe content or cell quantity. The R^2^ value of each regression line was used to evaluate linearity. Data are expressed as the mean±standard deviation from at least three independent experiments (technical replicates). All statistical analysis and data visualization was performed using Microsoft Excel.

## Acknowledgments

We gratefully acknowledge Dr. Adnan Bibic for his valuable assistance with the MRI experiments.

## Funding

This work was funded by grants from the National Institutes of Health (UH2/UH3 EB028904 (JWMB), S10 OD026740 (JWMB) and the Maryland Stem Cell Research Fund MSCRFD-5416 (JWMB) and MSCRFL-6270 (ASZ). As this manuscript is the result of funding in whole or in part by the National Institutes of Health (NIH), it is subject to the NIH Public Access Policy. Through acceptance of this federal funding, NIH has been given a right to make this manuscript publicly available in PubMed Central upon the Official Date of Publication, as defined by NIH.

## Author contributions

Conceptualization: JWMB, ASZ, SK, PW

Methodology: JWMB, ASZ, SK, PW

Investigation: ASZ, SK, CC, KS, AI, PW, JWMB

Visualization: ASZ, SK, CC, KS, AI, PW, JWMB

Funding acquisition: JWMB, ASZ

Project administration: JWMB

Supervision: ASZ, JWMB

Writing – original draft: ASZ, JWMB

Writing – review & editing: ASZ, SK, CC, KS, AI, PW, JWMB

## Competing interests

J.W.M.B. is a shareholder of SuperBranche. This arrangement has been reviewed and approved by Johns Hopkins University in accordance with its conflict-of-interest policies. Piotr Walczak is a founder and holds equity in IntraArt, LLC and Ti-com, LLC. Other co-authors have no competing interest to disclose.

## Data and materials availability

All data are available in the main text.

## Notes

### Summary of Updates

This version corresponds to the original manuscript submitted in 2025 and is being uploaded to comply with the journal preprint policy which allows only the initial submission (prior to editorial input and peer review) to be posted on a preprint server.

